# Sublethal exposure to the insecticide chlorantraniliprole does not alter antipredator behaviour in wild field crickets

**DOI:** 10.64898/2026.01.09.698581

**Authors:** Emily Gilford, Miranda Johnstone, Mark Pitt, Rolando Rodríguez-Muñoz, Chris Bass, Tom Tregenza, Bram Kuijper

## Abstract

Pesticide exposure has been proposed as a major driver of global insect declines. Sublethal effects occur when non-fatal pesticide exposure disrupts behaviour or physiology in ways that may reduce survival or reproductive success. However, most evidence for sublethal effects comes from laboratory studies under controlled conditions. We tested whether exposure to a sublethal dose of chlorantraniliprole, a widely used diamide insecticide known for its long environmental persistence, affects antipredator escape behaviours in wild field crickets (*Gryllus campestris*). Adult crickets were treated with either pesticide or a sham control. They then underwent behavioural tests in the wild. Following a simulated predator attack, we measured two key behaviours: escape speed and subsequent latency to re-emerge from their burrow.

We found no consistent effects of pesticide exposure on either behaviour. Re-emergence latency increased over successive days across both groups, while escape speed did not change. Re-emergence latency showed greater individual variability than escape speed. This likely reflects its role as a risk-assessment behaviour, which is more context-dependent than escape speed. Our findings challenge the assumption, largely derived from laboratory assays, that sublethal pesticide exposure universally impairs insect behaviour. Overall, our results underscore the need to integrate environmental context when evaluating pesticide impacts on insect behaviour.

## Introduction

Natural systems are increasingly shaped by anthropogenic disturbance, exposing wild animals to a range of novel and recurrent challenges. Insects, which underpin terrestrial ecosystems, are experiencing alarming declines, with over 40 percent of species threatened with extinction globally due to agricultural intensification, agrochemical pollution, and climate change (Sánchez-Bayo and Wyckhuys, 2019). Pesticides are now routinely detected on non-target insects, even in conservation habitats and semi-natural landscapes, far from areas of direct application, through processes such as spray drift, runoff, and atmospheric deposition (Albaseer et al., 2025; Brühl et al., 2021; Quandahor et al., 2024; Wan et al., 2025). These pervasive residues highlight that pesticide exposure is widespread, persistent, and rarely limited to target pest species.

Non-target insects perform essential ecosystem services, including pollination, natural pest control, and nutrient cycling; all of which can be disrupted by pesticide contamination (Albaseer et al., 2025; Goulson, 2013). Field surveys have documented the presence of pesticide mixtures on pollinators and predatory insects in agricultural margins and protected areas, suggesting that biodiversity losses are not explained solely by habitat conversion, but also by chemical drift and chronic contamination (Brühl et al., 2021; Sánchez-Bayo and Wyckhuys, 2019). Consequently, concern is growing not only about the lethal effects of pesticide use, but also about potential sublethal consequences for behaviour, physiology, and reproduction in wild insect populations.

While acute mortality remains the primary measure in pesticide risk assessment, there is increasing recognition that survival alone does not capture the ecological impact of exposure, particularly when measured in the laboratory. This has prompted growing interest in alternative measures of sublethal effects of pesticides which may reveal impairment in insects long before death. Sublethal pesticide exposure can alter physiology or behaviour by impairing movement, reproduction, or sensory perception, ultimately influencing fitness and population persistence (Gandara et al., 2024; Guedes et al., 2025) and has been proposed as a major driver of global insect biodiversity loss (van der Sluijs, 2020).

Despite the prevalence of pesticide drift and contamination, the vast majority of research on sublethal pesticide effects has been conducted in controlled laboratory environments, often on model taxa and under simplified conditions. Such work provides critical mechanistic insight, but fails to capture the ecological complexity of wild populations, which are exposed to variable doses and multiple interacting stressors that can alter both pesticide toxicity and behavioural responses (Brühl et al., 2021; Kaunisto et al., 2016; Slos and Stoks, 2008; Wan et al., 2025). For instance, pesticide effects can interact strongly with environmental variables such as temperature and humidity, with elevated temperatures often amplifying toxicity and sublethal effects, and lower humidity enhancing insect susceptibility to pesticides by increasing metabolic and desiccation stress (Tsaganou et al., 2021; Verheyen and Stoks, 2019). Understanding how sublethal pesticide exposure affects the behaviour of wild insects is particularly urgent, as declines are increasingly reported in landscapes with little or no direct pesticide application (Albaseer et al., 2025; Brühl et al., 2021; Sánchez-Bayo and Wyckhuys, 2019). Sublethal effects may also vary over time as individuals recover from, compensate for, or accumulate exposure impacts (Batool et al., 2024; Siviter et al., 2018).

Because of this, it is important to assess behaviour at multiple time points following exposure. Field studies are therefore needed to test how low-level exposures alter ecologically relevant behaviours. Only a few examples exist, and the range of insect groups and pesticide classes tested so far remains narrow, as most existing data concern older chemical groups such as pyrethroids, organophosphates and neonicotinoids and a limited set of model taxa (Biondi et al., 2013; Desneux et al., 2007; Guedes et al., 2025). Expanding this evidence base is essential for linking laboratory-derived mechanisms to behavioural and demographic outcomes in natural systems. Here we develop a field assay to assess the impact of pesticides in a wild population of insects.

In the context of natural populations, movement related traits, such as flight, walking, or burrowing, are particularly informative for understanding sublethal pesticide effects as they link directly to ecological performance. Mobility determines how insects locate food and find mates, and even subtle changes can have cascading effects on survival and fitness. Moreover, movement lies at the heart of antipredator strategies, as the ability to detect, evaluate and respond to threats is highly consequential to survival. Changes in mobility may therefore compromise risk-avoidance capacity and alter trade-offs between predator evasion and other activities such as foraging and reproduction. Escape behaviours are valuable indicators, as they integrate sensory, motor and motivational processes. Impaired escape may make individuals either overly responsive and wasteful of energy, or less responsive and vulnerable, depending on how pesticides disrupt neuromuscular and sensory processing (Adamo, 2012; Gandara et al., 2024; Guedes et al., 2025). Laboratory experiments and recent syntheses show that sublethal insecticide exposure can impair escape responses across a range of aquatic and terrestrial organisms (Ågerstrand et al., 2020; Bertram et al., 2022; Kavallieratos et al., 2024). This suggests that sublethal pesticide effects on fitness may be mediated through alterations in escape behaviours in response to threats and stressors. However, studies under natural conditions are essential to properly gauge the effects of pesticides on escape behaviours.

To address this gap, we combined a laboratory dose-response bioassay with a field experiment on wild field crickets. We first conducted a laboratory bioassay on the closely related field cricket *Gryllus bimaculatus* to characterise the dose-response relationship and identify a sublethal chlorantraniliprole concentration. We then applied this dose to wild field crickets (*G. campestris*) in northern Spain to test whether sublethal exposure alters antipredator escape behaviours. *G. campestris* is a well-studied orthopteran whose ecology and individual life histories have been monitored in long-term field populations, making it an ideal model for examining pesticide effects in natural environments (Rodríguez-Muñoz et al., 2025, 2019, 2010). Crickets exhibit a range of antipredator responses that are easily quantified in the field, including fleeing into burrows and the timing of subsequent re-emergence. We focussed on two behaviours with distinct ecological roles: escape speed, representing locomotor performance during fleeing, and re-emergence latency, reflecting post-disturbance risk assessment. Both behaviours were elicited using a simulated predator attack; a vibration caused by dropping a ball down a tube which landed close to the burrow entrance, which reliably triggers a flee response in this species (Gilford et al., (in press); Li et al., (in press), 2025).

We tested whether acute, sublethal chlorantraniliprole exposure influenced these escape behaviours and whether any effects persisted across two consecutive days of testing. Diamides such as chlorantraniliprole represent a rapidly expanding, multi-billion dollar class of insecticides that are increasingly replacing neonicotinoids in agriculture (Jeanguenat, 2013). Acting through ryanodine receptor modulation, diamides disrupt muscle function in target pests, while showing relatively low toxicity to vertebrates and other beneficial insects (Cordova et al., 2006; Nauen, 2006). Yet, despite the widespread adoption of diamides, little is known about their sublethal or behavioural effects on non-target species.

Chlorantraniliprole, one of the most widely used diamides in Europe, dominates insecticide applications in UK orchards (AFBI, 2022; Jeanguenat, 2013; Ridley et al., 2020) and is marketed as a reduced-risk, “bee-safe” alternative to neonicotinoids (Larson et al., 2013). However, emerging research supports subtle locomotor and reproductive impairments in bees and moths following exposure (Henry et al., 2012; Zhang et al., 2022). Chlorantraniliprole is also used extensively to control orthopteran pests such as rangeland grasshoppers and Mormon crickets *(Anabrus simplex*) (USDA APHIS, 2008), yet its sublethal effects on these groups, and other non-target orthopterans, remain poorly understood.

By integrating ecologically relevant behavioural measures with a free-living field population, our study provides one of the first assessments of how sublethal pesticide exposure shapes antipredator responses in wild insects. Based on laboratory findings that chlorantraniliprole disrupts muscle function in invertebrates (Cordova et al., 2006; Nauen, 2006), we predicted that sublethal exposure would impair antipredator escape behaviour in crickets by reducing locomotor performance and altering risk assessment. Specifically, we expected dosed crickets to exhibit slower escape speeds and longer re-emergence latencies compared to sham dosed controls. If sublethal effects accumulate or persist, we further predicted stronger behavioural impairments on the second day of testing. Conversely, if crickets rapidly recover following acute exposure, treatment effects would be transient or absent.

## Methods

### Laboratory bioassay for dose selection

To select a realistic sublethal chlorantraniliprole dose for use in the field, we ran a preparatory laboratory bioassay on the closely related Mediterranean field cricket, *Gryllus bimaculatus*, which can produce fertile hybrids with *G. campestris* (Tyler et al., 2013; Veen et al., 2013) and is commercially available as reptile food. Crickets were kept in a controlled environment at 24°C before and after dosing. We weighed technical grade chlorantraniliprole (PESTANAL® analytical standard; CAS No. 500008-45-7, Sigma-Aldrich, Supelco) to 0.01 mg accuracy (A&D HR-100A analytical balance, A&D Company Ltd., Japan) and dissolved it in 100% acetone to create a series of dose concentrations. Using a Burkhard hand micro applicator (Burkard Manufacturing Co. Ltd., Rickmansworth, UK), we administered a 2 μL drop of either solvent only (sham control) or one of seven chlorantraniliprole concentrations to the dorsal thorax of 240 adult male crickets (30 individuals per concentration). Cricket mortality was recorded at 24, 48, 72 and 96 hours after dosing. Here, mortality refers to the proportion of individuals found dead at each observation time; no time-to-event or survival analysis was conducted.

Test concentrations were 0 (acetone control), 1, 10, 35, 50, 100, 200 and 400 mg L^-1^ chlorantraniliprole. The 35 mg L^-1^ dose was calculated from the recommended field application rate of the formulated product Coragen^®^ (see Supplementary Methods), and the remaining concentrations were chosen to span a broad range around this value to allow us to fit a dose-response curve. This bioassay was designed to identify a concentration that produced low but non-zero mortality over the testing period, which we could then use operationally as a “sublethal” dose for the field experiment on *G. campestris* (see Results for justification).

### Natural study system

Our study site is a meadow in northern Spain, in which a population of the field cricket *G. campestris* has been observed by the “Wild Crickets” project since 2006 (see www.wildcrickets.org) (Tregenza et al., 2022). *G. campestris* are flightless, and spend most of their lives around self-dug burrows, which serve as refuges from predators throughout development and during winter diapause (Vrenozi and Uchman, 2020). Crickets typically forage and sun-bask near burrow entrances, retreating underground in response to predators, thermal extremes, or rain (Gardner et al., 2024). Nymphs emerge in spring, become more active as they develop, and reach adulthood in April–May (Rodríguez-Muñoz et al., 2011). The species is sensitive to substrate-borne vibration (Niemelä et al., 2015), and previous work has shown that vibrational stimuli reliably trigger rapid flight into the burrow, validating this response as an ecologically relevant antipredator behaviour (Gilford et al., (in press); Li et al., (in press), 2025).

We conducted field experiments in spring 2025, using *G. campestris* nymphs already present in the meadow from the 24th of April to the 9^th^ of May. In March, we searched the meadow for cricket burrows, which were identified with a uniquely numbered flag. We installed infra-red HD video cameras over 140 of the burrows, recording activity 24 hours a day, allowing behaviours and developmental stages of crickets to be tracked. In mid-April, crickets began emerging as adults. Upon observing adults with our monitoring system, crickets were caught two days post-emergence into their adult form to allow for pronotum hardening to facilitate tag adhesion, ensuring that all individuals were treated and tested at a consistent early adult stage. When caught, crickets were weighed (± 0.01 g), photographed, provided with a treatment (pesticide/sham) and marked with an acrylic tag glued to the pronotum to facilitate individual identification.

### Capture and dosing protocol

To examine whether sublethal exposure to chlorantraniliprole affects escape behaviours in wild *G. campestris*, newly emerged adults of both sexes were captured in the field and randomly assigned to receive either a topical dose of chlorantraniliprole (35 mg L^-1^, see bioassay results below) or a sham acetone treatment. Chlorantraniliprole dilutions were prepared using powdered chlorantraniliprole (PESTANAL® analytical standard; CAS No. 500008-45-7; Sigma-Aldrich, Supelco), weighed to the nearest 0.1 mg with an analytical balance (Mettler Toledo AG245; Mettler Toledo, Switzerland), and diluted to 35 mg L^-1^ in 100% acetone. A total of 294 adult crickets were treated: control individuals (*n* = 146) received a topical sham dose of 2 μL acetone, while pesticide-treated individuals (*n* = 148) received 2 μL of the chlorantraniliprole solution. For the present study, a subset of 59 individuals was randomly selected for behavioural assays (32 pesticide treated: 16 females, 16 males; 27 controls: 12 females, 15 males). Post-dosing, we returned crickets to the same burrows from which they were captured. Previous work in this population has shown that crickets handled and temporarily moved in this way resume normal activity on return to their burrows, with no detectable adverse effects (Rodríguez-Muñoz et al., 2025, 2019, 2010).

### Behavioural assay and stimulus delivery

We used a single-trial antipredator assay that simulates a predator attack via a standardized vibrational cue, following established protocols (Gilford et al., (in press); Li et al., (in press), 2025). We dropped a 42 mm, 9 g cork ball vertically through a 50 cm tube mounted on a tripod, landing approximately 10 cm in front of the focal cricket burrow. A GoPro camera (2.7K resolution, 240 fps) was positioned directly above the burrow to record each trial, with a 30 cm ruler placed next to the burrow entrance to provide a spatial scale for distance and speed measurements. Prior to each trial, we measured the body temperature of the focal individual using an infrared thermometer (Testo 830-T4), as escape performance in ectotherms is known to be temperature-dependent (Li et al., 2025).

Behavioural testing began one day after pesticide or sham dosing, allowing for capture recovery and pesticide absorption. As adult crickets emerged on different dates, dosing and subsequent test days were staggered across individuals. Consequently, “Day 1” and “Day 2” post-treatment refer to time since dosing, rather than to the same calendar days. We tested crickets once per day, with a maximum of two consecutive test days per individual if they remained present at a burrow. Of the total sample, 24 individuals were tested on Day 1 only, 2 on Day 2 only and 33 on both days. Burrow occupation was tracked using infra-red cameras, positioned over 140 of the burrows.

### Video analysis

We extracted escape speed from video recordings using ‘Kinovea’ (https://www.kinovea.org/download.html), which allows frame-by-frame analysis, with each frame representing 1/240^th^ of a second. We calculated escape speed over the first 1.5 cm of movement, as this provides a standardised estimate of initial response intensity before acceleration introduces variation. All crickets in this study fled at least 1.5 cm, making this a reliable and comparable segment across individuals. This approach follows previous experiments at the same field site, which used the same vibrational stimulus and found 1.5 cm to be sufficient for capturing early escape behaviour while minimising variation due to turning angle or distance to refuge (Li et al., 2025). The variables extracted from the video were; (1) the video frame in which the cricket’s first obvious movement was detected, which we denote as the response frame (*r_f_*), (2) the video frame in which the stressor-related cue was released (release frame), (3) the frame in which the cricket reached a distance of 1.5cm relative to its original position (*t_1_*) and (4) the frame that contained the maximum distance fled. Parameters were used to calculate escape speed (m/s). We calculated the time taken for a cricket to cover 1.5 cm (*t_x_)* using *t_x_* = (*t_1_ − r_f_*)*(1/240)). This was then calculated across the 1.5 cm segment to calculate the escape speed in meters per second (*f_s_*); *f_s_* = 0.015 / *t_x_*.

### Data Analysis

We fitted a multivariate Bayesian model to test whether escape speed and re-emergence latency differed between treatment groups (pesticide vs sham) and whether these behaviours were affected by days post-dosing (1 vs 2 days). The model included two response variables (1) escape speed and (2) re-emergence latency (log-transformed to improve normality). Fixed effects were treatment, days since treatment, sex, and mean-centred temperature, with individual ID (tag) included as a random intercept to account for repeated measures. Residual correlation between responses was estimated.

To assess whether pesticide effects varied with time since exposure, we compared models with and without a treatment × day interaction using leave-one-out cross-validation (LOO). Models were fitted in brms (v2.21.0) (Bürkner, 2017) under R (v.4.3.2)(R Core Team, 2023) using the rstan backend (Stan v2.32.2). We used default brms priors (flat priors on fixed effects and weakly informative priors on intercepts, variance components, and residual correlations) and confirmed via sensitivity checks that inferences were robust to alternative prior specifications (see Supplementary Methods). Gaussian likelihoods were used for both traits, following inspection of residual distributions and posterior predictive density-overlay checks comparing observed and model-generated distributions, and because alternative distributions did not improve model fit. Parameters were estimated using four MCMC chains (2000 iterations each; 1000 warmup), and convergence was assessed using R-hat statistics (with values close to 1 indicating good mixing) and effective sample sizes (ESS > 1000).

Posterior predictive checks stratified by days post-treatment verified that the models captured behavioural distributions well. LOO comparisons showed no improvement in predictive performance when the interaction was included (Table S1); therefore, we present results from the additive model.

### Ethics

This study was approved by the University of Exeter Research Ethics Panel (approval number: 9802182). A total of 59 individuals were used in this experiment across 92 observations and were caught and given either a pesticide or sham treatment. We carefully selected pesticide treatments based on prior laboratory work to ensure sublethal pesticide exposure levels and minimise mortality. All handling procedures were refined to minimise stress, and individuals were returned to their original burrows immediately following treatment and tagging. We subsequently monitored all crickets in the wild throughout their natural lives. While pesticide exposure may have caused behavioural or physiological changes, no procedures were performed that would be expected to cause pain, injury, or lasting distress.

## Results

### Laboratory dose-response bioassay

In insecticide bioassays conducted in the laboratory, mortality of *G. bimaculatus* increased with both chlorantraniliprole concentration and time since dosing (Fig 1). Based on mortality rates across the 96-hour period, we selected 35 mg L^-1^ as the concentration for the field experiment. In the laboratory bioassay, the 35 mg L^-1^ dose caused minimal but measurable mortality over 96 hours, so we treated it as an operationally sublethal exposure. The same topical dosing method and 2 μL application volume used in the laboratory bioassay were then applied to G. campestris individuals in the field experiment.

**Figure 1.**
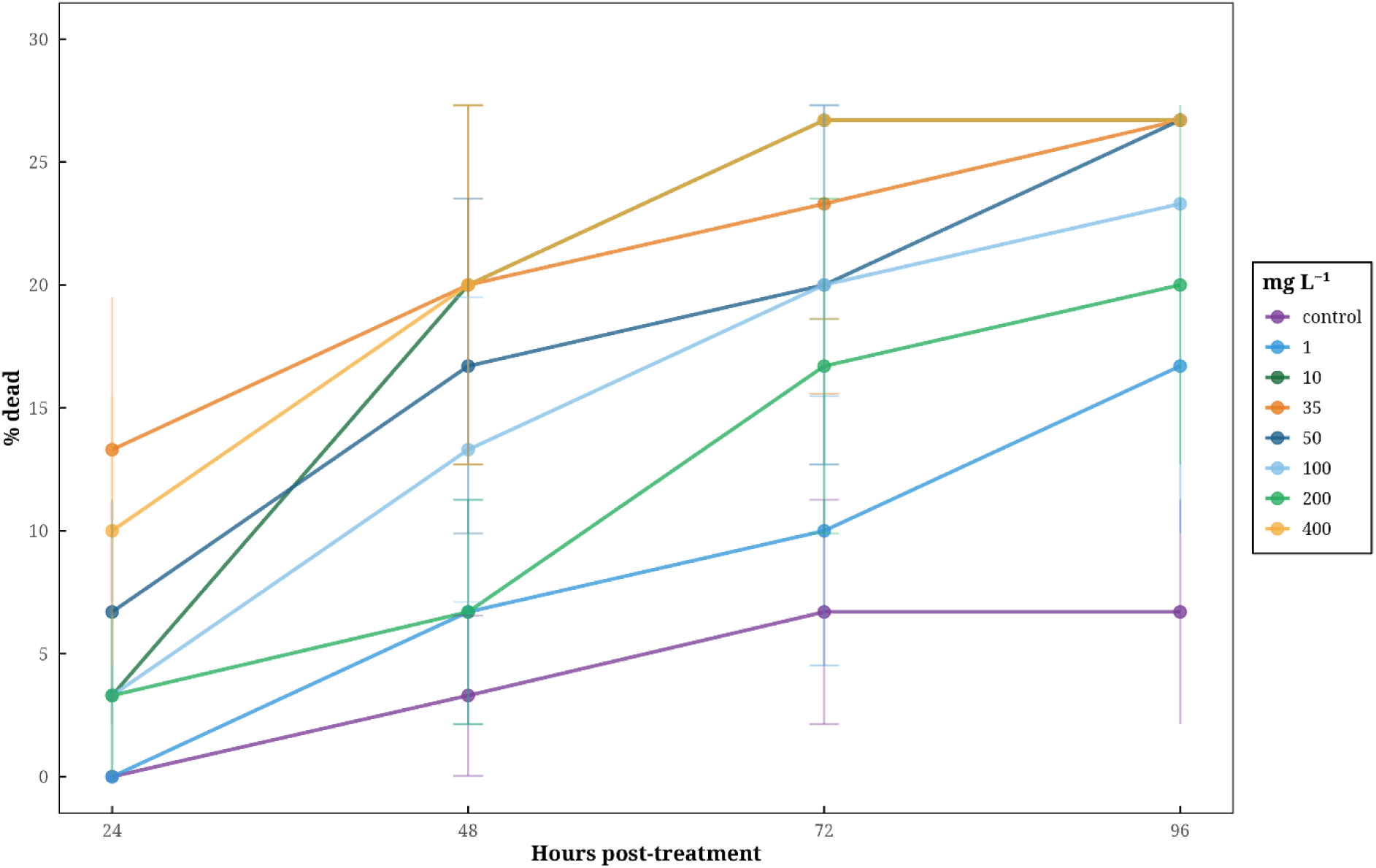
Laboratory bioassay results showing mortality rates of G. bimaculatus across a range of chlorantraniliprole concentrations and exposure durations. Percentage mortality was recorded across 24-96 hours post-treatment for an acetone control and seven pesticide exposure doses. Error bars represent standard errors. The selected sublethal dose for the field study (35 mg/L-1) is shown in dark orange, reflecting a dose causing minimal but measurable mortality over time. These results demonstrate dose- and time-dependent pesticide toxicity, supporting the assessment of behavioural impairments at one- and two-days post-treatment in the field experiment.

### Do pesticides alter escape speed and re-emergence latency?

Crickets exposed to the pesticide showed no statistically credible difference in escape speed compared to sham-dosed individuals (*β*_treatment:sham_= 0.00, 95% CI [−0.07, 0.08]; Fig 2B, Table S2). On the original scale, the estimated mean difference between groups was small (Sham − Dosed: median = 0.003 m s⁻¹, 95% CrI [−0.065, 0.075]). In addition, there was no difference in re-emergence latency between pesticide dosed and sham controlled individuals, as the confidence interval included zero (*β*_treatment:sham_= −0.24, 95% CI [−0.65, 0.19]; Fig 2A, Table S2); back-transformed to seconds, the estimated group difference was -8.5 seconds (Sham − Dosed; 95% CrI [−24.3, 6.9]), which was small relative to natural variation and not credibly different from zero. A summary of all fixed effect estimates is shown in Figure 4.

**Figure 2.**
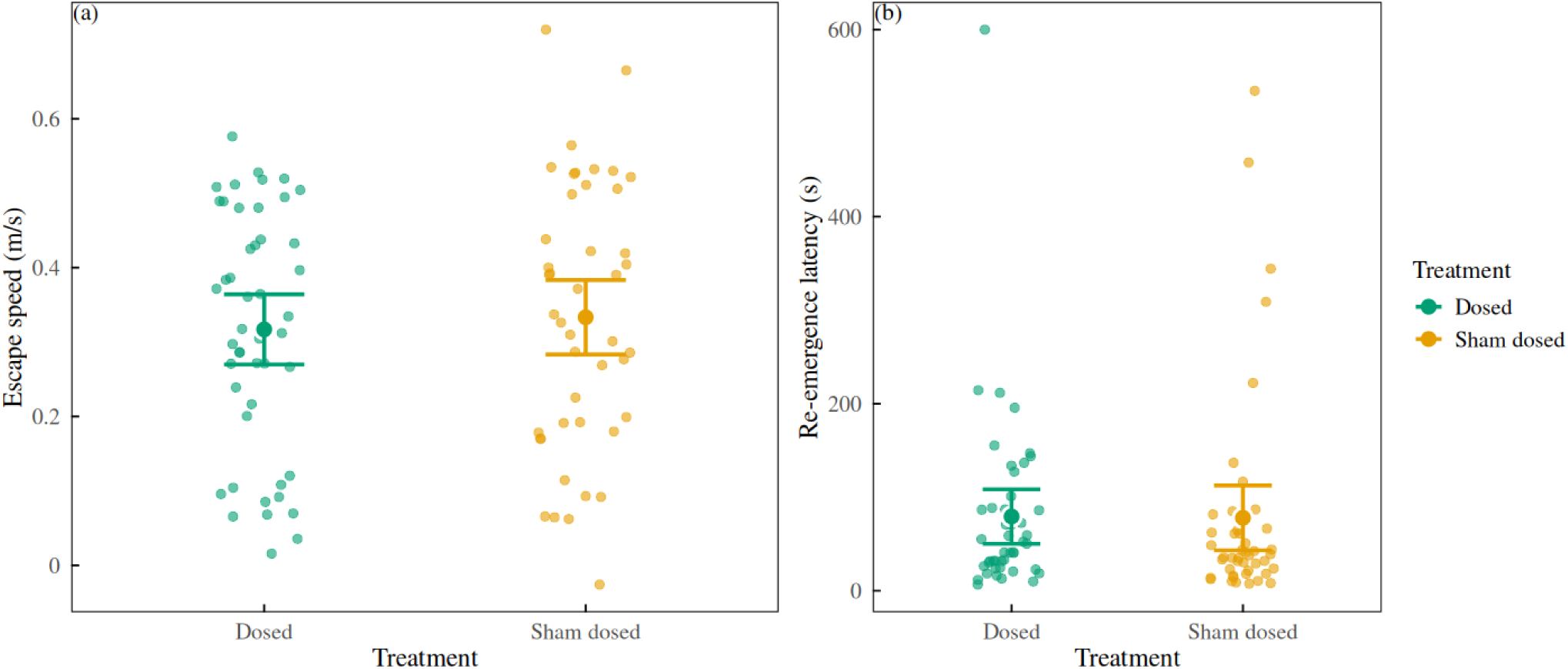
Effects of pesticide exposure on escape speed and re-emergence latency in adult crickets. (A) Escape speed (m/s) and (B) re-emergence latency (s) of crickets either dosed with pesticide (green) or sham-dosed (orange). Points represent individuals. Large points with error bars show posterior means and 95% credible intervals.

### Does time since treatment influence behaviour?

Posterior predictive checks stratified by days post-treatment indicated good model fit (Fig S1), with the models accurately capturing the distributions of both escape speed and log re-emergence latency for Day 1 (*n* = 57) and Day 2 (*n* = 35), despite the smaller sample size on Day 2. Crickets tested on Day 2 showed longer re-emergence latencies (*β*_post-treat:2_ = 0.45, 95% CI [0.08, 0.82], Table S2), corresponding to an increase of approximately 18 seconds on the original scale (median; 95% CrI [3, 40]), whereas escape speed did not differ by day (*β*_post-treat:2_ = −0.03, 95% CI [−0.10, 0.04], Table S2) (Fig 3).

**Figure 3.**
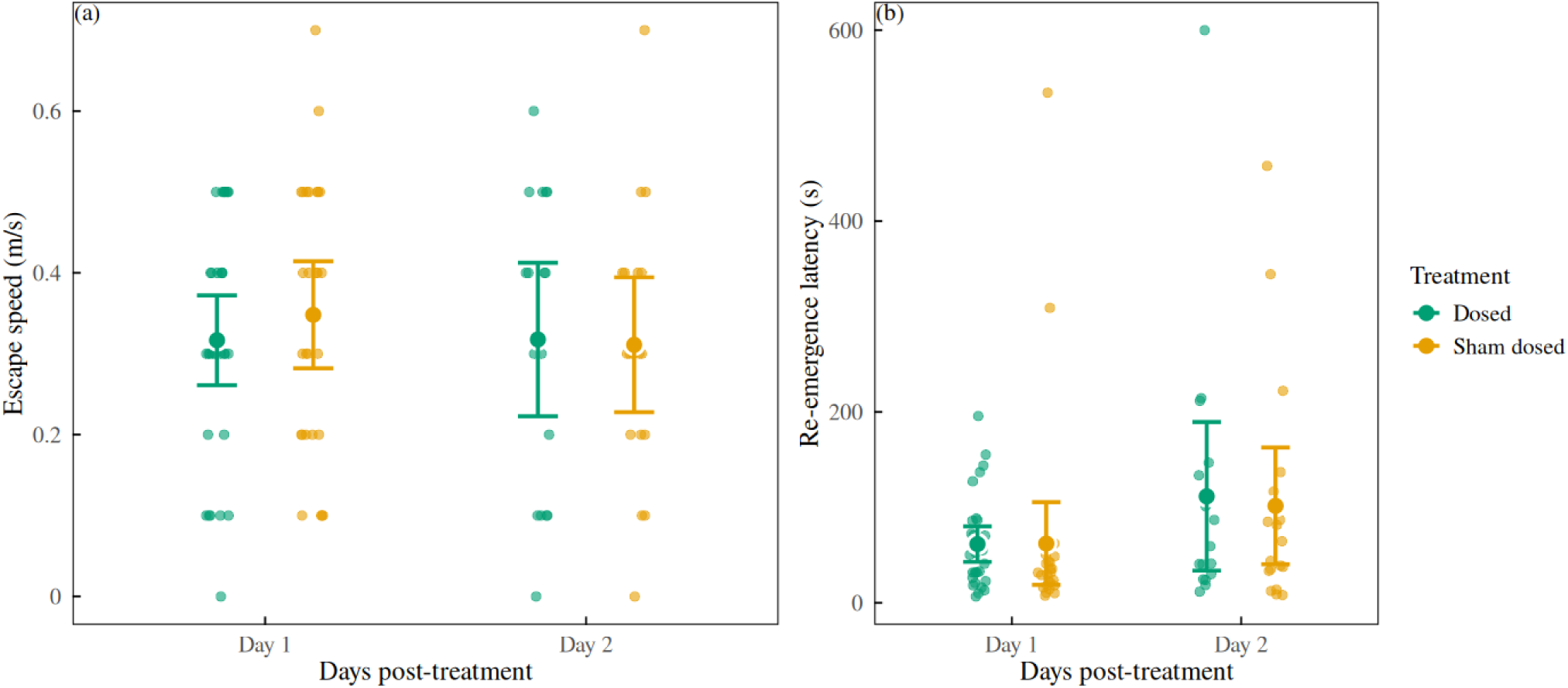
Effects of time since treatment on escape speed and re-emergence latency. Posterior mean (A) escape speed (B) and re-emergence latency for crickets dosed with pesticide (green) or sham dosed (orange), measured 1 and/or 2 days after treatment. Points represent individuals. Larger dots and error bars show the posterior mean and associated 95% credible intervals for each group.

**Figure 4.**
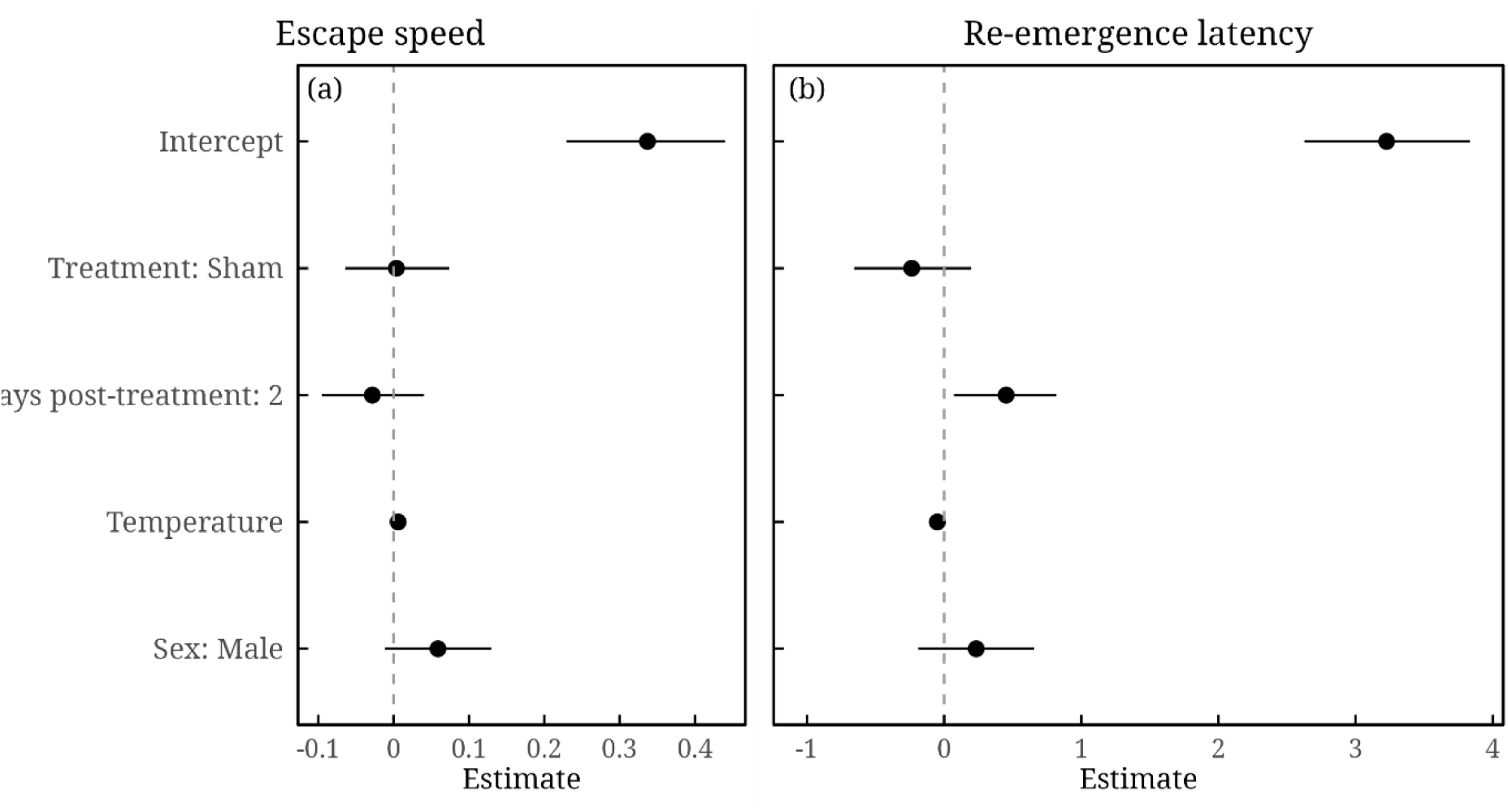
Forest plot of fixed-effect estimates for escape speed and re-emergence latency. Posterior mean estimates (black dots) and 95% credible intervals (horizontal lines) for the fixed effects: treatment (sham-dosed), day post-treatment (Day 2), temperature, and sex (male), shown for (A) escape speed and (B) re-emergence latency. The vertical dashed line at zero represents no effect.

### Do other factors affect escape behaviours?

In addition to the effect of pesticides on escape behaviour, we also explored the effects of temperature and sex. Temperature had a small but credible effect on both escape behaviours. Individuals tested at higher temperatures emerged more quickly (*β*_temp_ = −0.05, 95% CI [−0.08, −0.02]; Table S2) and showed slightly increased escape speeds (*β*_temp_ = 0.01, 95% CI [0.00, 0.01]; Table S2).

Sex showed no credible effect on re-emergence latency (*β*_sex:male_ = 0.25, 95% CI [−0.20, 0.66], Table S2) or escape speed (*β*_sex:male_ = 0.06, 95% CI [−0.01, 0.13]; Table S2).

### Consistency across individuals

Including individual identity (tag) as a random effect captured within and between-individual variability in baseline behavioural responses (Dingemanse and Dochtermann, 2013).

Posterior estimates for the standard deviation of the random intercepts were larger for re-emergence latency (sd = 0.47, 95% CI [0.10, 0.75], Table S2) than for escape speed (sd = 0.04, 95% CI [0.00, 0.10], Table S2), indicating greater between-individual variation in re-emergence latency.

## Discussion

### Do pesticides alter re-emergence latency and escape speed?

Our field experiment found no statistically credible effects of sublethal chlorantraniliprole exposure on re-emergence latency or escape speed in wild field crickets. These results challenge the assumption, largely derived from laboratory assays, that sublethal pesticide exposure universally impairs insect behaviour under field conditions. They align with growing evidence that behavioural effects of pesticides are context-dependent, varying across species, doses, and environmental complexity (Adamo, 2012; Ågerstrand et al., 2020; Bertram et al., 2022).

This result contrasts with many laboratory findings in other insects reporting sublethal pesticide-induced reductions in movement or escape responses (Delpuech et al., 2005; Stanley et al., 2015; Suchail et al., 2001), although several studies and meta-analyses have also reported weak or inconsistent behavioural impairment under certain conditions, highlighting that sublethal effects can be highly context dependent (Ågerstrand et al., 2020; Bertram et al., 2022; Melvin and Wilson, 2013). The absence of strong acute effects in our study may reflect intrinsic compensatory capacity in wild crickets, such as physiological or behavioural mechanisms that help individuals to maintain performance after low-level exposure. It may also reflect environmental buffering, where natural microhabitat conditions such as temperature variation, shelter, or access to resources, mitigate the realised toxicity of pesticides compared to simplified laboratory environments (Bro-Jørgensen et al., 2019; Gressel, 2011). Alternatively, sublethal effects may require more prolonged or repeated exposure to manifest (Sanchez-Bayo and Goka, 2014; Siviter et al., 2018).

Re-emergence latency increased modestly from Day 1 to Day 2 across both treatment groups, likely reflecting short-term behavioural plasticity rather than pesticide effects. This interpretation is supported by the absence of a treatment x day interaction, with both sham-and pesticide exposed individuals showing similar increases in emergence latency. Moreover, previous work at the same field site shows that repeated disturbance reliably leads to elevated re-emergence latencies in wild crickets (Gilford et al., (in press)), consistent with a plastic, experience-dependent response. This suggests that latency is a flexible, risk-sensitive trait influenced by recent experience, whereas escape speed appears more constrained and physiologically stable. Minor effects of temperature and sex were consistent with previous work showing that environmental conditions and individual state modulate antipredator behaviour (Li et al., 2025). Together, these results indicate that natural variation and repeated disturbance exert stronger influences on escape behaviour than acute, residue-level pesticide exposure.

### Implications, limitations and future directions

Despite modest or null effects, and acknowledging the inherent limitations of a short-term, single-dose field experiment, this study provides, to our knowledge, the first field-based test of sublethal chlorantraniliprole effects on antipredator behaviour in a wild insect population. Against a backdrop of rising diamide use and ongoing insect declines, behavioural experiments in the field are essential for validating laboratory-based hazard predictions and understanding how behavioural plasticity and environmental variability interact under real-world conditions (Cinel et al., 2020; Sánchez-Bayo and Wyckhuys, 2019).

Our findings suggest that acute, sublethal chlorantraniliprole exposure does not reliably impair core escape behaviours. Instead, most variation in both escape speed and re-emergence latency was explained by non-chemical factors, including temperature, sex, prior disturbance and between individual differences, each of which had larger and clearer effects than pesticide exposure. Re-emergence latency in particular proved more flexible and risk-sensitive than escape speed, responding readily to recent experience. This indicates that short-term behavioural plasticity and common environmental stressors, such as repeated disturbance, predator cues or thermal conditions, occupy a more influential position in shaping behavioural variation in the wild than residue-level pesticide exposure. In our case, increases in re-emergence latency are more plausibly attributed to normal life experience (e.g. sensitisation to repeated disturbance) than to pesticide itself, consistent with recent findings that repeated non-lethal stressors elicit sensitisation rather than suppression of antipredator behaviour (Gilford et al., (in press)). We therefore conclude that acute, residue-level exposure to chlorantraniliprole at the dose applied here did not measurably alter escape speed or re-emergence latency in this wild population. This finding does not preclude effects at higher doses, under repeated or prolonged exposure, or on other behavioural or physiological traits, or through interactions with naturally occurring environmental stressors, all of which remain important directions for future research.

Sublethal effects may, however, emerge after longer or repeated exposure. In long-term laboratory studies, behavioural and physiological disruptions were observed days or weeks after dosing, particularly when pesticide exposure occurs alongside additional environmental stressors such as elevated temperatures or repeated disturbance, and in some species even across generations (Batool et al., 2024; Sanchez-Bayo and Goka, 2014; Siviter et al., 2018). Ideally, future studies on wild populations would therefore focus on long-term and multigenerational approaches. However, as in the current study, this is not always feasible due to the effort required to track individuals over extensive periods of time.

A key limitation of the present study is that it focused on a conservative, residue-level exposure rather than direct-spray or repeated dosing scenarios. The chlorantraniliprole concentration used here represented a conservative, residue-level exposure, lower than direct field deposition (i.e. the amount an insect would receive if struck directly by spray droplets during application). A 2 μL topical application at this concentration delivers approximately 70 ng of active ingredient, around ninefold lower than the estimated dose a cricket would receive under direct spray deposition (≈651 ng, based on a mean exposed surface area of 1.86 cm²).

Spray-drift and off-crop deposition studies show that only a small fraction of the applied pesticide reaches field margins and adjacent vegetation (Grella et al., 2017; Longley and Sotherton, 1997), indicating that the exposure used here is consistent with drift-derived contamination of non-target organisms. Field surveys similarly report pesticide residues on non-target insects and vegetation in agricultural margins and conservation habitats (Albaseer et al., 2025; Brühl et al., 2021). This likely reflects the most common exposure pathway for non-target insects, particularly ground-dwelling species such as *G. campestris* that spend much of their time in burrows and are therefore rarely struck directly by spray, but instead encounter residues carried by spray drift or atmospheric deposition onto vegetation and soil surfaces (Albaseer et al., 2025; Brühl et al., 2021; Solé et al., 2024). Our findings therefore do not rule out behavioural impairment under higher, repeated, or cumulative exposures, but indicate that single, residue-level exposure is not a dominant driver of the behavioural dynamics measured here. Future work could test higher, near-application concentrations to evaluate potential behavioural impairment in directly exposed individuals and better bracket the range of ecologically relevant doses.

Understanding the physiological pathways underlying behavioural change, such as modulation of octopamine and related neurochemical systems (Adamo, 2017), will be important for clarifying how exposure translates into altered risk responses. Mechanistic perspectives can reveal constraints on plasticity, help explain species-specific differences in vulnerability, and identify the points at which sublethal stressors begin to compromise neuromuscular or sensory processes. Such insights are essential for linking individual behaviour to population-level consequences and for improving predictions of how reduced-risk pesticides affect wild insects.

In summary, our findings support calls for ecologically realistic behavioural measures in pesticide risk assessment and continued emphasis on field studies that capture natural behavioural variability and individual lability in response to pesticide exposure. By revealing how field-relevant behaviours respond to environmental context and stressors often missed or misrepresented in laboratory assays, this work advocates expanding field-based behavioural assessments as essential complements to traditional ecotoxicology.

## Conclusion

To our knowledge, this is the first field test of sublethal chlorantraniliprole effects on antipredator behaviour in a wild insect. We found no strong or consistent impairments in escape performance following pesticide exposure at the dose employed, although re-emergence latency exhibited temporal sensitivity and between-individual variability, supporting its potential as an early and responsive indicator of sublethal stress in ecological settings. In an era of accelerating chemical use and biodiversity decline, linking behavioural plasticity to environmental toxicology remains crucial. By testing predictions largely derived from laboratory studies in a natural population, our results highlight that the behavioural consequences of sublethal pesticide exposure may be weaker or more context-dependent in the field than often assumed, and further studies are warranted. This work contributes to the growing recognition that behavioural traits, particularly those related to decision-making and antipredator performance, offer functionally relevant windows into how sublethal stressors may influence organismal fitness and ecological interactions.

## Notes

### Competing Interest Statement

The authors have declared no competing interest.

### Funding

- Natural Environment Research Council: NE/S007504/1

