## Supplementary Materials for "Sublethal exposure to the insecticide chlorantraniliprole does not alter antipredator behaviour in wild field crickets"

### Methods

#### Dosage selection calculations

We selected a working concentration of 35 mg L^-1^ chlorantraniliprole for the field experiment. This solution was applied topically at 2 μL per individual, delivering approximately 70 ng of active ingredient per cricket. This dose was chosen to represent a realistic sublethal exposure. For comparison, the recommended field application rate of chlorantraniliprole for orchard pest control is approximately 35 g active ingredient (a.i.) per hectare, equivalent to about 0.35μg cm^-2^ under uniform deposition. According to the Coragen^®^ MAPP 14930 product label (Du Pont (UK) Limited 2016), the formulation contains 200 g L⁻¹ chlorantraniliprole and is recommended at 175 mL ha⁻¹ in apples and pears, corresponding to 35 g a.i. ha⁻¹. An adult field cricket with an exposed dorsal area of roughly 1.86cm^2^ (mean of 24 measured individuals), could therefore receive approximately 0.65μg (651ng) of active ingredient under direct spray, whereas our applied dose was around nine times lower. The 35 mg L⁻¹ dose also produced only minimal mortality over 96 hours in our preparatory laboratory bioassay, confirming that it was appropriate for testing sublethal behavioural effects in the field.

#### Data Analysis

Analysis was conducted in R (v4.3.2) (R Core Team 2023). Data and an analysis script are available as supplementary materials. The script performs data import, cleaning, and preprocessing, including converting re-emergence latencies from mm:ss format to seconds and mean-centring temperature values to facilitate interpretation of model coefficients. The script also produces figures visualising behavioural patterns and posterior estimates, which are included in the manuscript and supplementary materials. Predictive calibration was additionally assessed using leave-one-out probability integral transform (LOO-PIT) ECDF diagnostics for each response; these showed no strong departures from uniformity, indicating adequate predictive calibration. Default brms priors were used, which was appropriate given the lack of strong prior information for treatment effects in this system. These comprised flat priors on fixed effects and weakly informative Student-t priors on intercepts, variance components, and residual correlations, providing regularisation without overpowering the likelihood. To assess prior sensitivity, we refit the model using more and less regularising priors; posterior estimates for the treatment and day effects were qualitatively unchanged (posterior means and 95% credible intervals were similar, and conclusions were identical). Posterior distributions showed reasonable behaviour and were not truncated at the bounds of the priors, supporting the adequacy of this approach. For ease of interpretation, we additionally report selected model-implied differences on the original response scales, calculated from posterior expected values and back-transformed where necessary.

Table S1

Model comparison using Leave-One-Out cross-validation (LOO) between the additive model (rsmodel_add) and the interaction model including treatment × days post-treatment (rsmodel_int).

| **Model LOO comparison** | *elpd_diff* | *se_diff* |
| --- | --- | --- |
| **rsmodel_add** | 0 | 0 |
| **rsmodel_int** | -3.0 | 0.80 |

The table reports the difference in expected log predictive density (elpd_diff) and its standard error (se_diff), relative to the best-performing model (set to zero). Differences in predictive performance were small (elpd_diff <4 and elpd_diff/se_diff <4) indicating no meaningful improvement in predictive accuracy when the interaction was included. We therefore retained the simpler additive model.

### Results

#### Tables

Table S2

Posterior summaries of the fixed effects from the Bayesian multivariate mixed-effects model (rsmodel_add) assessing the effects of chlorantraniliprole exposure on antipredator behaviours in wild field crickets.

| **Fixed effects** | *Parameter* | *Estimate* | *SE* | *Lower 95% CI* | *Upper 95% CI* | *Rhat* |
| --- | --- | --- | --- | --- | --- | --- |
| **Escape speed** |  |  |  |  |  |  |
|  | **Intercept** | 0.34 | 0.05 | 0.23 | 0.44 | 1.00 |
|  | Treatment: Sham | 0.00 | 0.04 | -0.07 | 0.08 | 1.00 |
|  | Days post-treatment: 2 | -0.03 | 0.03 | -0.10 | 0.04 | 1.00 |
|  | Sex: Male | 0.06 | 0.03 | -0.01 | 0.13 | 1.00 |
|  | Temperature | 0.01 | 0.00 | 0.00 | 0.01 | 1.00 |
| **Re-emergence latency (log)** |  |  |  |  |  |  |
|  | **Intercept** | 3.23 | 0.30 | 2.64 | 3.82 | 1.00 |
|  | Treatment: Sham | -0.24 | 0.21 | -0.65 | 0.19 | 1.00 |
|  | Days post-treatment: 2 | 0.45 | 0.19 | 0.08 | 0.82 | 1.00 |
|  | Sex: Male | 0.25 | 0.22 | -0.20 | 0.66 | 1.00 |
|  | Temperature | -0.05 | 0.02 | -0.08 | -0.02 | 1.00 |

The model included escape speed and re-emergence latency (log-transformed) as response variables. Predictors were pesticide treatment (sham/dosed), days post-treatment (1/2), sex (male/female), and temperature. Posterior mean estimates, standard errors, and 95% credible intervals are provided. Rhat values indicate model convergence, with values close to 1.00 suggesting satisfactory convergence. Intercepts represent the estimated baseline value when all predictors are at their reference levels: sham treatment, one day post-treatment, female sex and mean-centered temperature.

Table S3

Posterior summaries of the group-level standard deviations (SDs) for the random intercepts in the Bayesian mixed-effects model (rsmodel_add).

| **Random effects** | *Parameter* | *Estimate (SD)* | *SE* | *Lower 95% CI* | *Upper 95% CI* |
| --- | --- | --- | --- | --- | --- |
| **Escape speed** | |  |  |  |  |
|  | **Tag (random intercept)** | 0.04 | 0.03 | 0.00 | 0.10 |
| **Re-emergence latency (log)** |  |  |  |  |  |
|  | **Tag (random intercept)** | 0.47 | 0.16 | 0.10 | 0.75 |

This model assesses individual-level variation in antipredator behaviours: escape speed and re-emergence latency (log-transformed). Random intercepts for individual identity (tag) were estimated for each behavioural response. The table reports posterior mean SDs, their standard errors (SE), and 95% credible intervals (CI). These values reflect between-individual differences in baseline behaviour. Intercepts represent the estimated baseline value when all predictors are at their reference levels: sham treatment, one day post-treatment, female sex, and mean-centered temperature.

#### Posterior Predictive Plots

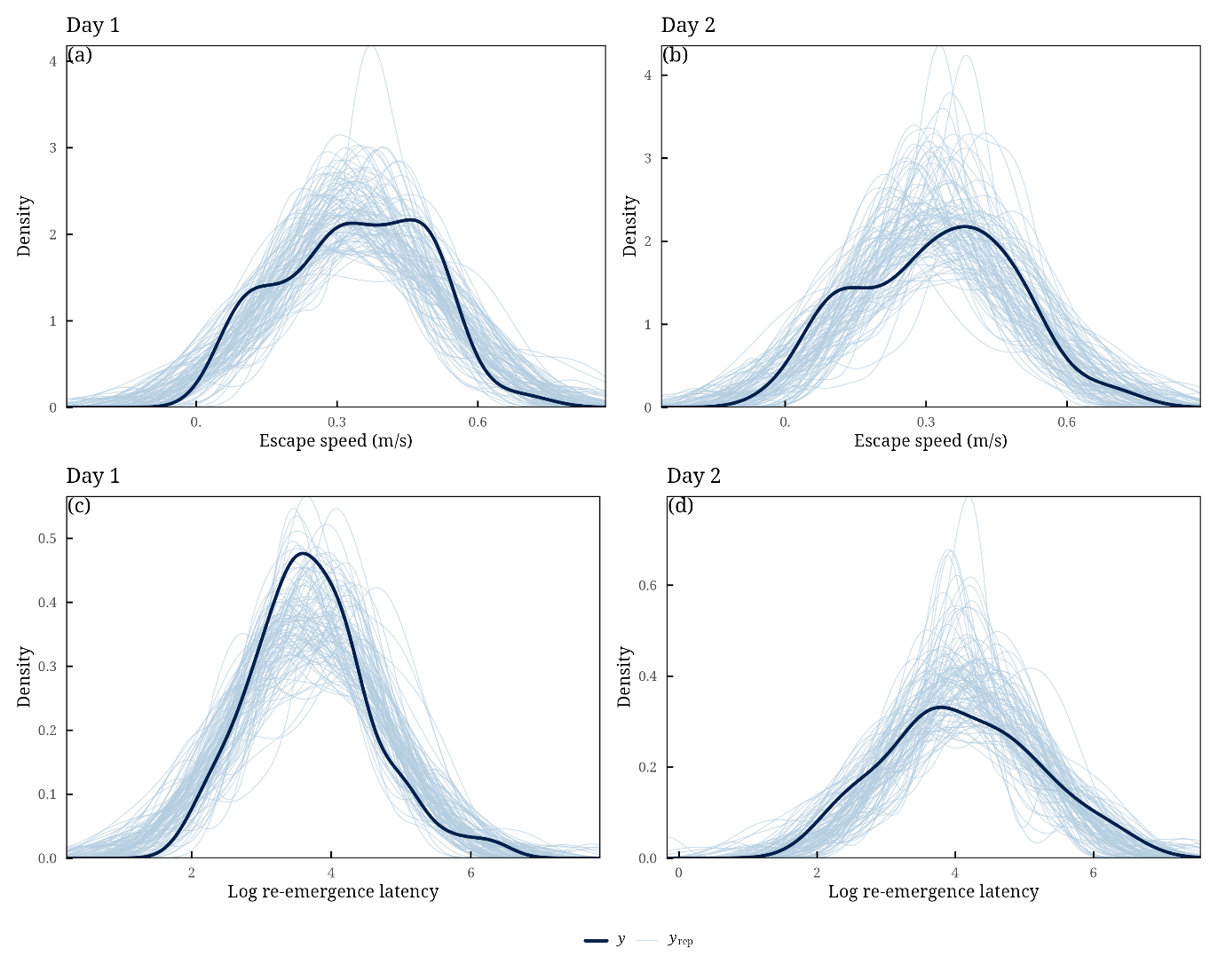
Figure S1 – Posterior predictive density plots for escape speed and log-transformed re-emergence latency across days post-treatment in the rsmodel_pestbd model. Plots show observed data distributions (dark lines) and 100 posterior draws (light lines) for: (A) escape speed on Day 1, (B) escape speed on Day 2, (C) log re-emergence latency on Day 1, and (D) log re-emergence latency on Day 2. The model captures the central tendencies and spreads of behavioural responses across both days, despite differences in sample size (Day 1 = 57, Day 2 = 35). These plots support the model’s ability to reproduce the observed data structure across the two time points

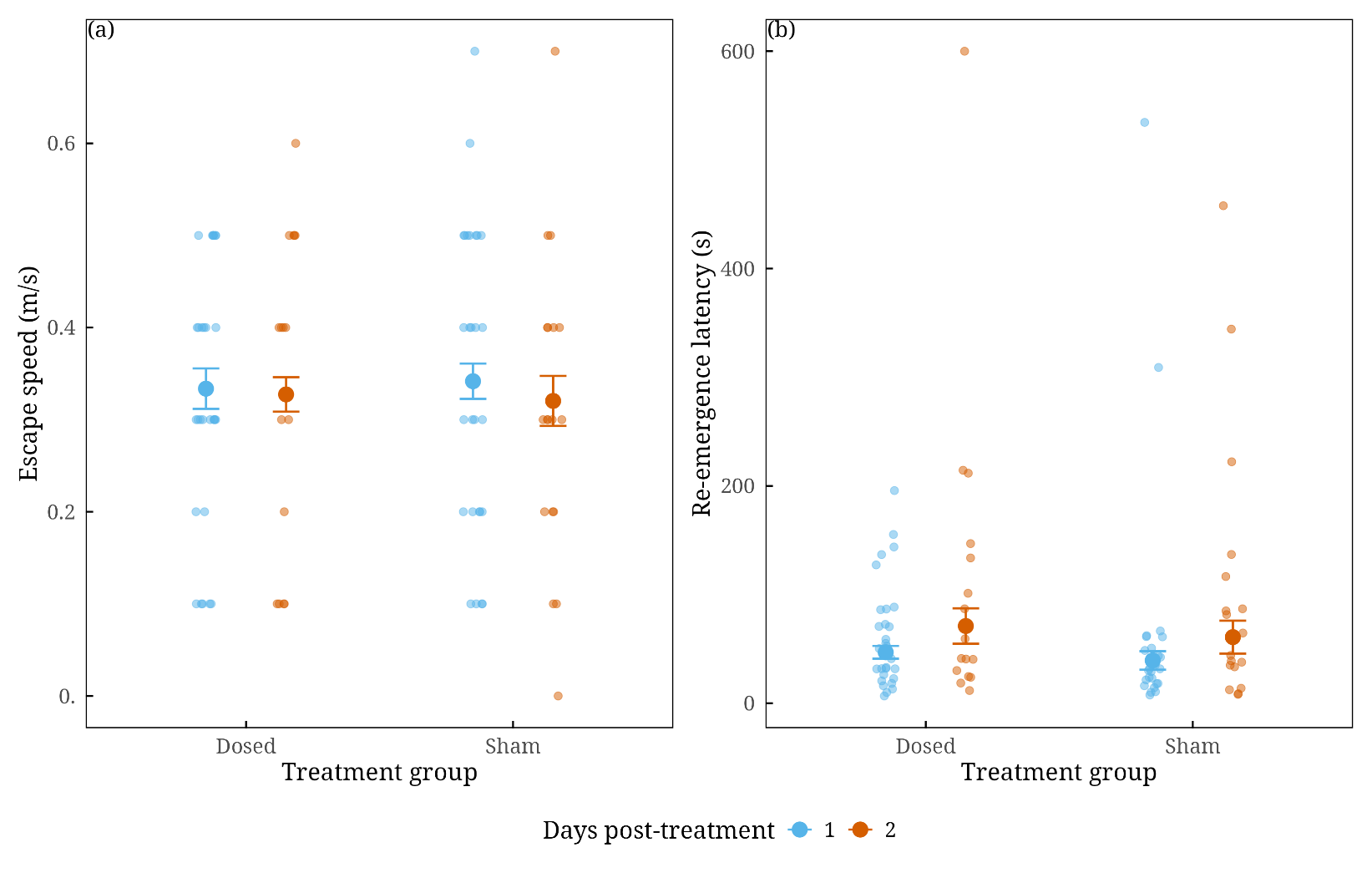

Figure S**2** – Posterior predictive plots, showing predicted behavioural responses from rsmodel_pestbd presented in Table S1 across two days post-treatment (Day 1 = blue; Day 2 = orange) for crickets dosed with either a sublethal pesticide treatment (Dosed) or a sham control (Sham). Large points represent posterior mean estimates of the fixed effects; vertical lines show 95% credible intervals. Jittered points indicate observed values used in the model. Plots show: (A) escape speed and (B) re-emergence latency. Predictions were generated with other covariates (temperature and sex) held constant at reference values (mean temperature and female sex) and exclude random effects. All values are shown on the original scale.
